# The proposed promiscuity value of an HLA can vary significantly depending on the source data used

**DOI:** 10.1101/2021.10.01.462828

**Authors:** Jordan Anaya, Alexander S. Baras

## Abstract

Immune checkpoint blockade, a form of immunotherapy, mobilizes a patient’s own immune system against cancer cells by releasing some of the natural brakes on T cells. Although our understanding of this process is evolving, it is thought that a patient response to immunotherapy requires tumor presentation of neoantigens to T cells and patients whose tumors present a wider array of neoantigens are more likely to derive benefit from immune checkpoint blockade^1–4^. Manczinger et al.^5^ recently reported findings that would appear contrarian to this notion in that they suggested patients with HLA alleles which bind more diverse peptides (higher promiscuity) are less likely to respond to immunotherapy. To estimate HLA promiscuity they looked at the HLA-peptide binding repertoires for class I alleles contained in the IEDB^6^, and obtained consistent results when performing robustness checks and subsequent analyses. Here we show that the proposed HLA promiscuity values can vary significantly across source data types and individual experiments.

---

The authors define promiscuity (*Pr*) as 1*/KL* where *KL* is the average Kullback–Leibler divergence (*D*_*KL*_) across each position, *y*, of a peptide:

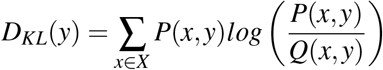

*P*(*x, y*) is the position-specific amino acid distribution observed in the HLA-peptide population and *Q*(*x, y*) is the position-specific background distribution. The authors included any peptide in the range of 8-12 amino acids which had either a positive HLA binding or T cell assay. They then limited themselves to alleles which had at least 400 associated peptides and used the peptides that can be generated from the human proteome for the background distribution.

We found this metric potentially interesting but had several concerns about how the source data used in its calculation could significantly affect the derived values. The data in the IEDB contains a variety of data source types ranging from HLA/MHC binding assays that test peptides predicted or known to bind to HLA alleles^7^ to mass spectrometry experiments of peptides found on the cell surface presented by HLA alleles^8^. In the case of mass spectrometry, using the proteome as the background distribution *Q*(*x, y*) is reasonable given that the antigen processing pathway acts on the proteome of the cells, albeit the peptide generation process is not completely random^9^. We would also note that differences in gene expression across cell types can be substantial; however, without a gene expression profile of some form for the cell line employed this cannot be modeled for. In contrast, we postulate that in the context of the HLA binding assay data the background distribution need not be modeled from the human proteome since the entirety of the tested peptides is explicitly known in these experimental settings. Even if an appropriate background distribution were to be used, the divergence from it would be expected to be much lower in binding assays as the *Q*(*x, y*) is often already lower in entropy compared to the human proteome. As a result, it’s difficult to understand how one might go about comparing *Pr* values from these two classes of experiments.

The issue of the different data types gave us enough concern that we performed a recent download of the IEDB database and, along with peptides from a recent large monoallelic dataset not yet in the IEDB^10^, then looked at the correlation of *Pr* values calculated with mass spectrometry as compared to HLA binding data, wherein we observed no correlation (Figure 1A). We next wondered what the correlation was between experiments which in theory should be more comparable, so we checked the correlations for various large mass spectrometry experiments^10–14^ (Figure 1B-G). In general there was better correlation between the allele *Pr* values, and this did not require both experiments to be monoallelic, but the scale of the values appeared to depend on the experiment. To investigate this we took 8 alleles which were common across three monoallelic data sets and plotted their values along with the values which would result from using all of the HLA and T cell assay data, which is the proposed protocol of Manczinger et al. (Figure 1H). The values vary wildly and it is clear that even among mass spectrometry experiments it is difficult to compare raw *Pr* values. The lack of correlation of *Pr* values derived from mass spectrometry versus HLA binding data was expected given the vastly different composition of these two data sources, but the lack of high correlation across independent monoallelic mass spectrometry data sets, along with the varying scales of the *Pr* values, indicates more validation needs to be done before these values are incorporated in prognostic and/or predictive models.

**Figure 1.**
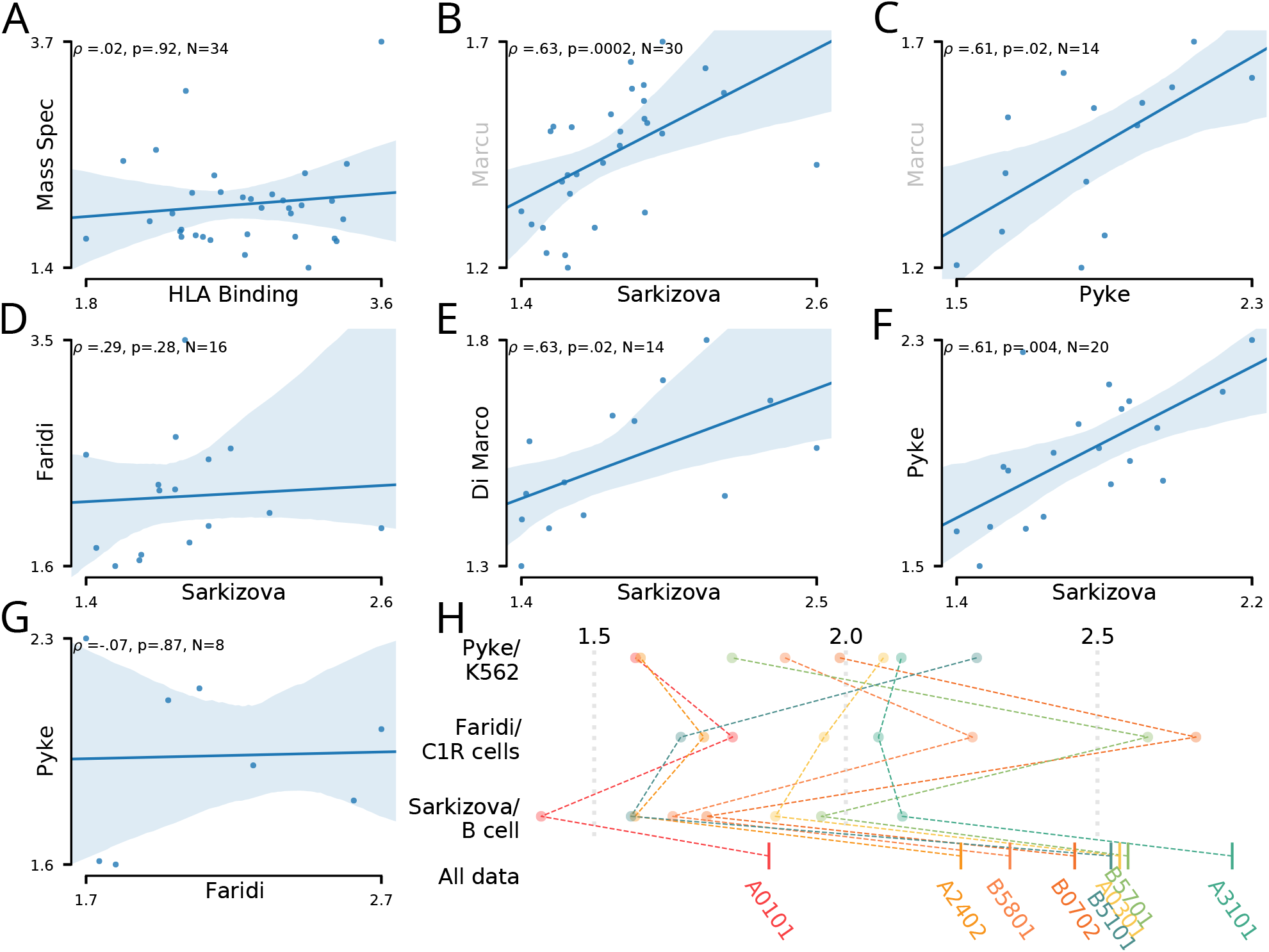
Experimental bias of promiscuity values. (A) Correlation of allele *Pr* values between HLA binding and mass spectrometry data. (B-G) Correlations of allele *Pr* values between mass spectrometry datasets. Monoallelic datasets are in black, with the sole polyallelic dataset in gray. (H) *Pr* values for 8 alleles which were present across three different monoallelic datasets, along with the value calculated with all data sources (including HLA and T cell assay data). Alleles are connected across references with dashed lines.

Although unlikely, it’s possible that the different alleles in the IEDB have data from enough different sources that any biases operate evenly across all alleles. Figure 2A shows the *Pr* values of Manczinger et al. along with the estimated proportions (see methods) of each data type. Not only do the different alleles have very different data sources, it’s clear that the alleles with higher *Pr* values (primarily HLA-A and HLA-B alleles) have a much larger proportion of HLA binding data. Mass spectrometry assays are becoming increasingly common to investigate the immunopeptidome, so we wondered how the data proportions have changed since the Mancinzger et al. publication and how that impacts the *Pr* values. As can be seen in Figure 2B, the mass spectrometry data type has significantly increased in its proportion. As might be expected given the overlap of the data used to calculate *Pr* originally and the available IEDB data at this point in time, a strong correlation (*ρ* = .84) was observed in Figure 2C. Interestingly, the addition of mass spectrometry data resulted in decreased *Pr* values, which is shown in (Figure 2D). As more and more mass spectrometry data is deposited into the IEDB, we expect that these values will continue to deviate from the original *Pr* values over time. We also checked that our code faithfully reproduces the values reported by Manczinger et al., as can be seen in the accompanying GitHub repository.

**Figure 2.**
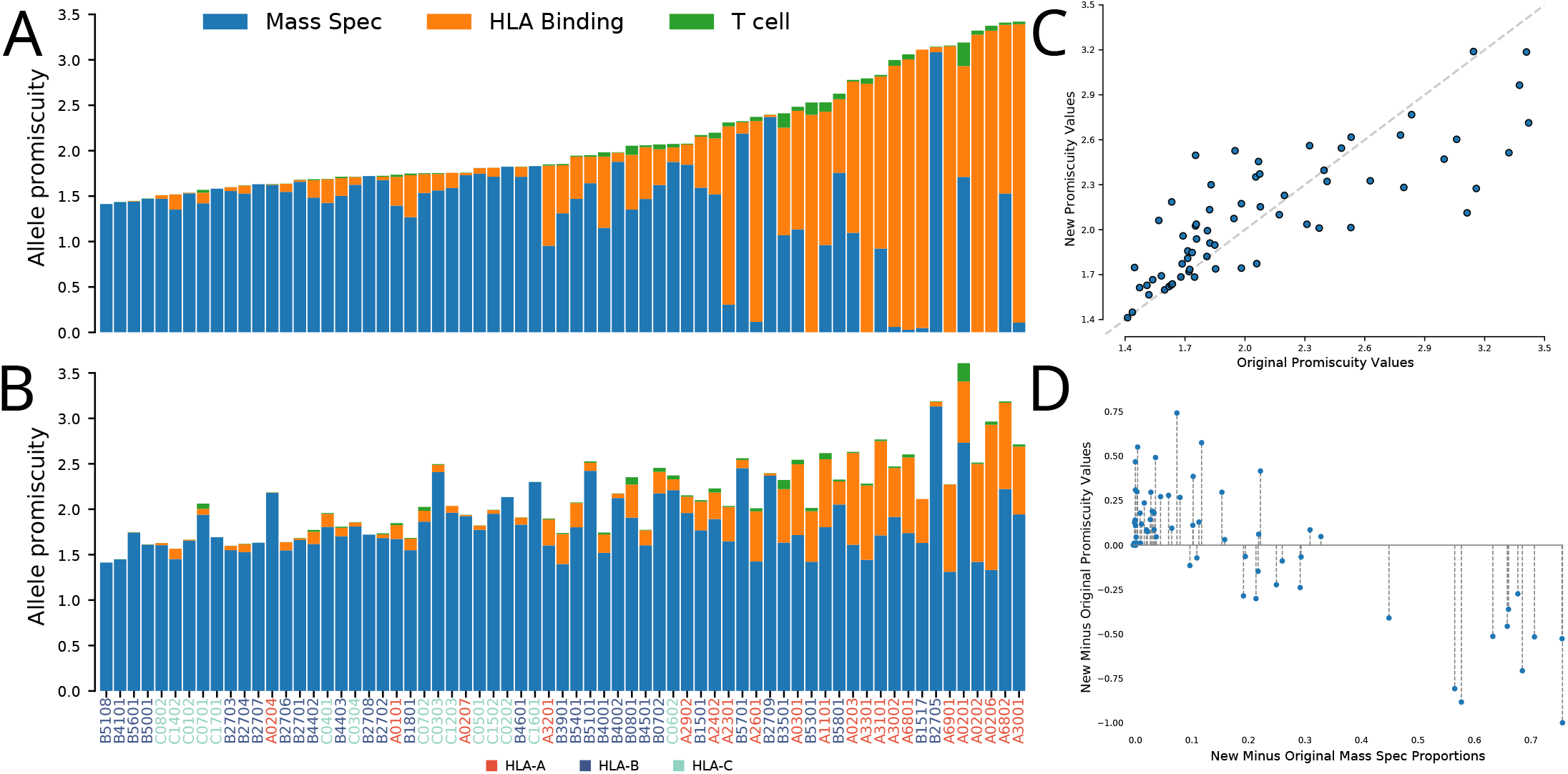
Assay composition of HLA class I alleles over time and the effect on *Pr* values. (A) The original *Pr* values of Manczinger et al., with the fill representing estimated proportions of the different data types. (B) *Pr* values calculated with a recent download of the IEDB database, and the fill representing proportions of the different data types. (C) The new *Pr* values plotted against the original *Pr* values. The line y=x is shown in gray. (D). The change in *Pr* values plotted against the change in mass spectrometry proportion.

While it’s encouraging that *Pr* values of mass spectrometry experiments correlate, it’s unclear how consistent the metric would need to be for it to see effective utilization. Theoretically the metric should be more accurate if data from multiple references are combined in its calculation, but given the different scales seen between references it’s unclear how *Pr* values should be combined. Despite the fact that clearly much work still needs to be done to understand how to best calculate this metric, Manczinger et al. obtained supporting results for their *Pr* values as a true metric of diversity. This is likely the case because it appears that the analyses are confounded by data overlap. The authors demonstrated that their *Pr* values correlate with peptide diversity in the Abelin et al. data^8^, but this is part of the data included in their *Pr* value derivations. They also used NetMHCpan to compare the predicted fractions of epitopes bound, however NetMHCpan is trained with IEDB data^15^. *Pr* values as currently calculated are largely driven by the relative decrease in entropy at anchor positions, and this potentially has biological implications; however, given the inconsistencies we’ve identified more work is needed to refine the calculation of HLA promiscuity as proposed by Manczinger et al.

## Availability of Data and Materials

The code provided by Manczinger et al. was converted from R to Python, and R was used to convert their. RData files into text files to be read by Python. Code for reproducing all results and figures of this manuscript is available at https://github.com/OmnesRes/hla_promiscuity, and has been archived at http://doi.org/10.5281/zenodo.5544924. From the database export (http://www.iedb.org/database_export_v3.php) of IEDB we downloaded “mhc_ligand_full” and “tcell_full_v3”. The peptides of Pyke et al. were extracted from their Supplemental Table 1, with peptides that contained modifications removed. To map the peptides provided by Manczinger et al. to references we matched the alleles and sequences to assays in the IEDB and then limited the references to those which had a date of 2018 or earlier since Manczinger et al. downloaded their data in January of 2019. Any technique in mhc_ligand_full which contained mass spectrometry in its name was considered a mass spectrometry data type and all others were considered HLA binding assays. Every reference in “tcell_full_v3” was considered a T cell experiment.

## Author Contributions

J.A. interpreted and analyzed data and prepared the manuscript. A.B.S interpreted data and prepared the manuscript.

## Competing interests

The authors have no competing interests.

## Funding

This research was supported by the Mark Foundation for Cancer Research (19-035-ASP), and the philanthropy of Susan Wojcicki and Dennis Troper in support of Computational Pathology at Johns Hopkins.

## Notes

http://doi.org/10.5281/zenodo.5544924

### Competing Interest Statement

The authors have declared no competing interest.

